# Amino acid stylization: a new approach for representing protein sequences and scoring protein alignments

**DOI:** 10.1101/270488

**Authors:** Stefano M. Marino

## Abstract

The amino acids (AA) stylization, introduced in this work, is a mean to obtain a qualitative and quantitative description of different AA; the goal is to allow, via a chemically meaningful representation, the analysis of protein similarity in multiple alignments, without any statistical and evolutionary information involved. The AA Stylization Matrix Method (aSMM) has been tested positively with different data sets, in line with most common statistically based approaches (e.g. BLOSUM62), while allowing the exploitation of some unique capabilities, e.g. to be rapidly configured for specific properties that can be (over-, or under-) weighted. Overall, applying *ad hoc* parameterization aSMM permits a tailored investigation to account for different types of substitutions in protein alignments.

## Introduction

Protein sequence similarity searching programs like Blastp or Fasta use scoring matrices to account for differences between residues in two or more aligned proteins [1-4]. Amino-acid changes can range from biochemically conservative, e.g., leucine to valine or glutamic acid to aspartic acid, to dramatically different, e.g., tryptophan to alanine [1]. The earliest amino-acid scoring matrices were based on amino acid properties or genetic code differences; instead, in the last decades amino-acid scoring matrices based on empirical measurements of amino-acid replacement frequencies (from sets of homologous sequences) became more and more used. Particularly, the BLOSUM family of substitution matrices (in particular the BLOSUM62 matrix) is currently the *de facto* standard in protein database searches and sequence alignments (e.g. it is the default choice, in the widely used web version of blastp at ncbi) [5,6]. The primary motivation behind this work comes as a consequence: to date, the work on statistically derived matrices has reached a relatively stable status (and matrices like Blosum62 are expected to remain essential for the foreseeable future) [1,6]; meanwhile, protein alignment scoring based on amino acid properties has been comparatively neglected, and the derivation of an efficient scheme remains an important challenge. In the following text, an approach for amino acid similarity scoring relying solely on amino acid properties description, through an innovative scheme (called amino acid stylization), is presented and discussed.

## Method

Amino acids, AA, share similarities and common features: while far from a continuous space (e.g. each amino acid being derived from another, by continuous additions or subtraction of one specific property), an opportune AA stylization scheme could permit a classification based on partial derivations (to represent, with approximation, AA diversity in an effective, albeit not strictly rigorous, manner). This was the goal of this work, and the stylization (with its underlying assumptions) represents the core of the innovation.

During the setup process, I developed and tested several alternative schemes, based on a manually curated, “human expertise” driven investigation. The pillars of the stylization, were:

1) side chains are represented as atoms in a grid; the representation is meant to be as fair as possible (but not strict), particularly in respect to size
2) functional groups and particularly non carbon atoms are considered of special importance
3) the stylization tries to capture the presence of other features, such as charge, aromaticity and carbon branching.

After preliminary analyses (i.e. on a range of alternative stylizations), a specific scheme was selected, as reported in Fig. 1.

**Figure 1.**
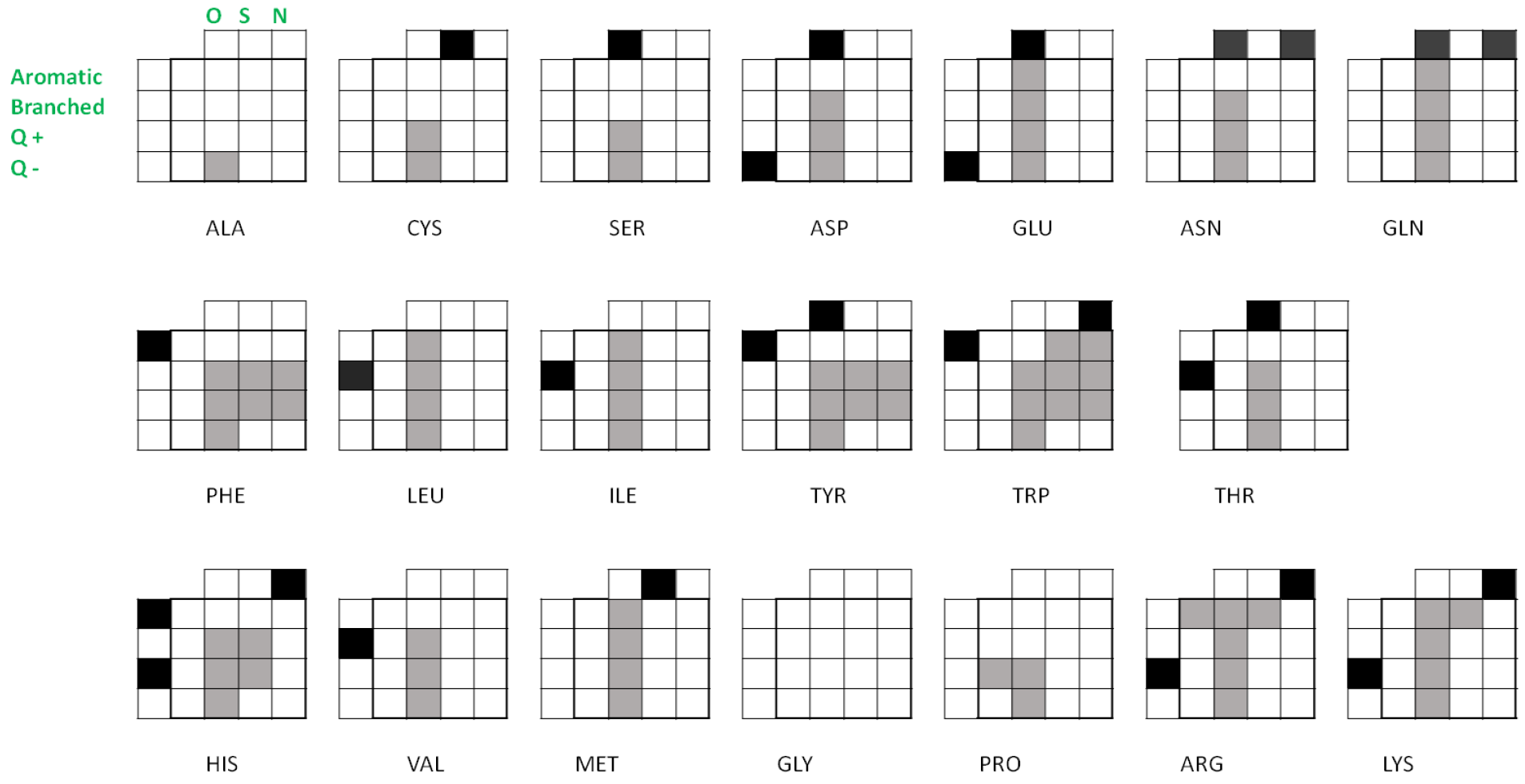
Amino acid stylization and features representation. First row and first column represent non carbon atoms and physicochemical features, respectively (see notation for Ala; O= oxygen, N=nitrogen, S=sulfur, Q+ = positive net charge, Q− = negative net charge). In the first column and first row (i.e. external layer, for features), a dark gray spot indicates positions with a full weight; Gly (special case) is scored with a default value(1, post assignment).

The example shows that for each amino acid, a 5×5 matrix composed of basically two regions is created: the 4×4 “inner core” region (inner section, from 2^nd^ row and from 2^nd^ column, representing the main non-hydrogen atoms of the side chains) and the “external” region, made of the first row and the first column (Fig. 1, first row includes the weights for non carbon atoms; first column the weights for some chemical/physical features: aromatic, branched, positive or negative charge). In terms of atomic representation, the main rationale of the stylization is: sidechain (non-hydrogen, nonH) atoms are represented in the “inner core” section (and treated equally, regarded of the atom type), while the presence of non-carbon atoms (e.g. oxygen, nitrogen, sulfur) is accounted in the first row; for example, delta position in Methionine is represented with an atom assignment in the inner section (e.g. as if it was a carbon) with a contemporary representation of sulfur in the external section (first row, positive weight for sulfur atom presence); in other words, all non-hydrogen sidechain atoms are treated equally in the inner section, while delegating the differences in functionalization to the external section.

This scheme was meant to have a visual component, and was meant to contain (for future developments) a specific color scheme; however, for the purpose of the present study, a grayscale stylization was developed, that could be directly implemented in matrices (one for each residue type).

As said before, the stylization includes the “inner core” of the matrix, and an “external” region for functional groups and features. Initially, all the non-empty positions in the matrix (squares in Fig. 1) are assigned a numerical value of 1 (i.e. nonH sidechain atoms=1, oxygen=sulfur=nitrogen=1, aromatic=branched=net_charge_positive=net_charge_negative=1). Then, a differential weighting is achieved by multiplying (via entrywise product, denoted with a° symbol in the following text) an amino acid specific matrix, e.g. M_i_, with a (5×5)weighting array, e.g. M_k_. The weighting array can specify for 8 differential weights, for any of the categories described.

After preliminary tests (inclusive of some optimization runs, see also later on the text), the default parameters of the derivation scheme in Fig. 1 were set as follows: w(inner_core_nonH_atoms)=1.0;w(oxygen)=w(nitrogen)=w(sulfur)=1.5;w(aromaticity)=w(branched)=w(positive charge)=w(negative charge)=1.0 An example of the “weighted” matrix for Serine, M_i_, could be:

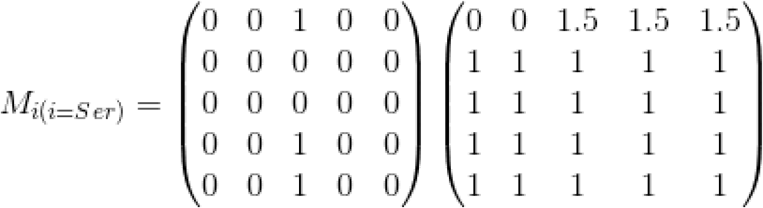

Generally, the score for each position in a two proteins sequence alignment is given by the absolute value of the difference between two matrices (|Ma – Mb|, where Ma is the weighted matrix for the amino acid x in sequence a, Ma=M(x) ° M_k_, and Mb is the same for sequence b), normalized by the sum of non zero positions (N) in both matrices.

In general, the score for the position *i*, S(*i*), can be expressed as

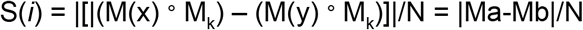

where, for a position *i* in the alignment, x is the aligned amino acid in sequence a, and y the corresponding amino acid in sequence b, M_k_ is the weighting matrix (previously defined, as well as the size normalization parameter, N).

Ultimately, the score of the alignment, is the sum of S(*i*) scores iterated for each aligned position, and divided by the alignment length.

Optimizations were run with all weights made to vary (independently) between 0.5 and 2.0 (with 0.5 intervals); the default scheme(all weights of 1.0, except oxygen, nitrogen, sulfur atoms, with a weight of 1.5,) was deliberated after rounds of optimization (of the weights distribution scheme, in parametric form) with reference datasets (made of externally generated multiple alignments, e.g. pfam and CDD); ultimately, the chosen default scheme was the most balanced in our tests, the one providing consistent performances in all the different scenarios tested. As a reference in the optimization, the program ClustalW was employed, with the scoring function, CW, derived as follows:

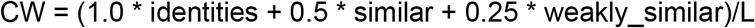

where similar are those positions marked by the symbol (“:”) and weakly_similar are those positions marked by the symbol (“.”), and L is the number of aligned positions.

After having defined its basic structure, the approach was further refined by introducing some “special” cases, for some peculiar amino acids: Gly, Pro, Cys. Indeed, with the stylization method presented, the description of the amino acids can be (with very little work) enriched with “custom” features; for example, in respect to the default scheme (see Fig. 1 and related discussion), small variants were introduced (Fig. 2): “special cases” were defined, for *(i)* Cys(fractional weight, 0.5x, for negative charge, at the light of its “close to neutral pH” pKa; a significant proportion of Cys, particularly if exposed, can exist, more or less transiently, in the thiolate form, [7]), *(ii)* Pro(fractional weights, 0.5x, to approximately represent its unique structural features, Fig. 2B); as mentioned in the legend of Fig. 1, Gly was already treated as a special case, with an extra matrix value of 1 (so that, for example, Gly − Ala = |0− Ala| + 1).To be noted, these modifications for the special cases, were introduced *a priori*, i.e. before testing their effect on the performances, mainly because these three adjustments were deemed as more opportune, fitting representations of these peculiar AA types. Two minor customization were also considered for Asn and Gln: to account for the “mild” (e.g. if compared with their close relatives, Asp and Glu) functionalization of their (side chain) functional groups, their carboxamide N and O atoms were fractionally weighted (0.5x, e.g. for the oxygen spot in the matrix, Asn is scored half the value of Asp). While biochemically grounded, these modifications provided overall minor effects in our tests (on average, for the tests in the Results section, less than 1% on the final score; naturally, the difference would become more significant, in alignments with unusually high number of Asn and Gln). For this study I opted not to use them (but it could be considered for future customizations or further refinement studies). Hereinafter, all values, numbers and stats reported in the Results section refers to the inclusion of the customized matrices for Cys and Pro in Fig. 2.

**Fig. 2.**
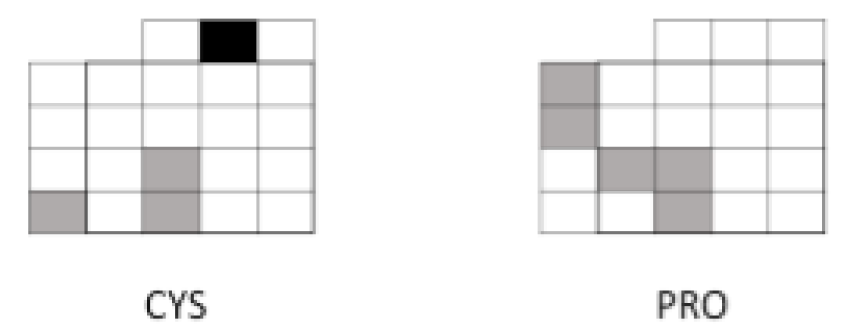
Customization for special cases. Cys and Pro present some fractional (0.5x) weights depicted in light grey.

An important mention goes for the treatment of Amino acid identity: by design AA identities are all treated equally (score contribution = 0), in aSMM with default option; this is not the case for most common matrices (e.g. Blosum, PAM)), which factors in also the different abundance (e.g. different probability of identity by chance for different AA types); in aSMM it is possible to introduce a dedicated scoring to account for this; in a preliminary study, I introduced a (very mild) scoring scheme of: −0.1 for Cys-Cys and Trp-Trp identities, −0.05 for Met-Met, His-His, Tyr-Tyr (and 0.0 for all other cases): the analysis revealed that the correlation between aSMM (with separate identity scoring) and CW and Blosum62 had a minor improvement (on average, 1%) if tested with Example 1 (pfam families, Results section). However, it may be important to consider the possibility to employ differential weights for identities in future developments. Last, but not least: gaps are accounted for, by default, with a score of +1. This is a conservative choice; however, gap weighting is an important parameters, that could require adjustments (e.g. more impact, for alignments of similar proteins, with few expected gaps), depending on the user needs. In the following section, all results refers to aSMM with default options (described before) for gaps and identity scoring.

## Results

To test the stylization approach introduced in the previous paragraphs I defined some alignments with *(i)* different protein families, and *(ii)* designed peptides with variable % id (from 38% to 87%, alignment length of 56 aa).

Overall, the scheme of Fig. 1 (aSMM, with the modifications of Fig. 2) returned consistent results, in line with two widespread evaluation methods tested (Clustal W based similarity scoring, CW, and Blosum62 similarity scoring). Moreover, by playing with the parameters, aSMM can change the results in a predictable way, providing a clearer picture in relation to user specific needs, i.e. favoring the conservation of specific functional features, such as aromaticity over electrostatics, or functional groups over size, etc.

### Example 1 – Comparative tests with different protein families

We used pre-generated alignments available in pfam (http://pfam.xfam.org/), for the following protein families: pfam16674, pfam15620, DUF1679, pfam00246 [8]. For the extended list of sequences employed, see Appendix. By analyzing the correlations between the scores of aSMM and ClustalW, for each alignment in the multiple alignment file, and then evaluating for each family the Pearson correlation (R^2^ coefficient) we obtained the following results (in brackets the values for the correlation between aSMM and BLOSUM62): R^2^[pfam16674] = 0.9345 (0.9388), R^2^[pfam15620] = 0.9191 (0.9115), R^2^[DUF1679] = 0.9590 (0.9630), R^2^[pfam00246] = 0.9260 (0.9339). These results highlight the significant agreement of aSMM with the two other methods: this is particularly remarkable at the light of the fact that while both CW and Blosum62 are derived from statistical analysis of amino acid frequency and evolution in natural proteins, aSMM is straightforwardly derived from a (chemically and structurally inspired) stylization, with no a priori knowledge of amino acid preferences in protein alignments.

### Example 2 – A closer look: method-specific differences and features

Considering the following input set of sequences:

~~~
>p5/1–56
PCTWDDFRGHLGGGNRKLVLIGWDEHEDMRLEFFFGNPLDKWWNYSEFERLLELLA
>p6/1–56
PCTHDKRVGHLGGGGGGGGGGGWNEHEDLPLEIFETNAQDKWWNYSDWSRAVELLA
>p9/1–56
CCCCCCCCCCCGGGGGGGGGGGWEEHEDMRLEIFEGNPLDKWWNYSEFERLLELLA
>p10/1–56
CCCCCCCCCCCGGGQHRLLIVGWEEHEDMRLEIFEGNPLDKWWNYSEFERLLELLA
~~~

These sequences were aligned (with clustalO, each pair aligned separately), scored with the three methods (CW, Blosum62, and aSMM), and sorted by similarity, as reported in Fig. 3 (from top to bottom, more similar to less similar; note that for aSMM lower scores stand for higher similarity, while the opposite is true for CW and Blosum62); the results show that, there is an overall consensus (indeed, all three methods individuate the best and the worst alignments); relative distance between different positions, however, was peculiar for each of the three methods, and reflect different takes on the evaluation of protein conservation (e.g. evolution driven for Blosum62, structural and chemical affinity for aSMM). Thus while one can confidently use different methods to detect the bigger trend, more subtle differences may require using different methods.

**Figure 3.**
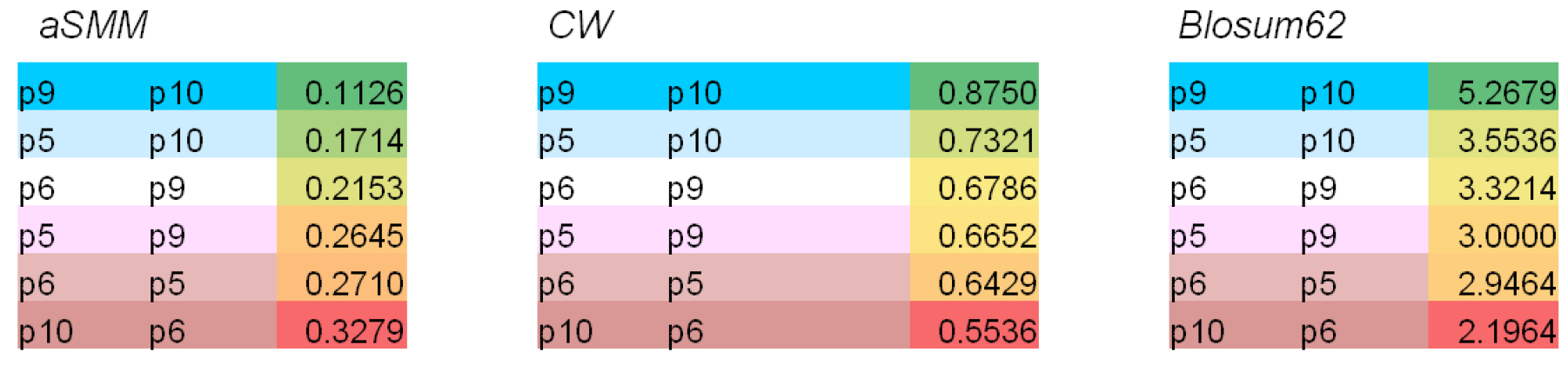
**Detailed view of the scoring and sorting** for each pairwise alignment in Example 2, for the three methods tested. Color codes (consistent between different cases in the figure) are used to facilitate the reader; color codes for scores: from green (best) to red (worse).

For what it regards this document, a first major highlight is the ability of such a different and independent approach (aSMM) to provide reliable results (for the general trend, in an overall agreement with Blosum62, but also with CW).

However, a major reason to develop (and to use) the method introduced here is the ability to tailor the analysis, and to customize the weight of different components and features.

An example of this application is reported below.

### Example 3 - Effects of weight schemes in aSMM scoring and alignment resolution

Considering the following input set of sequences:

~~~
>p6
GGGWNEHEDLPLEIFETNAQDKWWNYSDWSRAVELLATASLIPHFILA
>p7
GAAWNDYDCAPLEIFEVQADERYWNYSDWSKGTCTLAVGSIVVHYILG

>p8
GGSWNEHECLPLEGFETNAQDKVVCSCRWSRAAEPLATAYLKHHHISS
~~~

The scores for each alignment, and for different scoring methods are reported in Table 1.In this case, aSMM is in agreement with Blosum62, while CW is not (however, the latter is in agreement with the scoring by amino acid identity, ID in Table 1); it is important to note, that if other common matrices are considered (for clarity, in Table 1 only a representative set of matrices tested were reported), the trend is a consensus with aSMM and Blosum62, in spite of the fact that amino acid identity is considerably higher for the alignment preferred by CW.

**Table 1.** Results of different scoring matrices for Example 3.

| *protein 1* | *protein 2* | *aSMM* | *%ID* | *CW* | *Blosum62* | *Risler* | *structure* | *PAM30* |
| --- | --- | --- | --- | --- | --- | --- | --- | --- |
| *p8* | *p6* | 0.2309 | **0.6458** | **0.6771** | 2.8750 | 1.3833 | 2.5625 | 2.3333 |
| *p7* | *p6* | **0.1937** | 0.4792 | 0.6458 | **3.0417** | **1.4875** | **2.7917** | **2.4583** |
| *p7* | *p8* | 0.3408 | 0.3333 | 0.5000 | 1.7500 | 1.1083 | 1.4167 | 0.1458 |

This example was designed on purpose to create a “borderline” situation, to highlight that similarity scoring methods (e.g. the reference standard Blosum62, but also the aSMM approach) can overrule decisions made only on amino acid identity. Furthermore, the aSMM scores reported here refer to the default scheme of parameters; with a different weighting scheme, e.g. in this case favoring aromaticity and net charge, the relative scorings in Table 1 can become more separated: for example, with relative weights of 2.0 (for aromaticity and for both positive and negative charge) the scores become: p8-p6 = 0.2728, p7-p6 = 0.2073, p7-p8 = 0.3925 (which represents a +35% increase in the average difference between the best alignment, p7-p6, and the other two; the difference can increase much further, e.g. up to +95%, with higher relative weights for aromaticity and charge, e.g. weight = 5.0). Thus, the aSMM scoring allows for the possibility to enhance, specifically for a chosen set of features, the relative scoring (indeed, with the weighting scheme discussed above, aSMM allows for the best separation in the example, in respect to any other method in Table 1), and this is a characteristic feature of the approach presented here, which can be useful in several situations, where comparisons of differential scoring can highlight specific properties of the alignments. Naturally, the user should be aware of possible consequences of (heavily) biasing the scoring system (i.e. with “strange” weighting schemes, alignments can deviate significantly from those obtained with the default scheme).

## Discussion

The amino acid stylization method (aSMM) presented here is an original and orthogonal addition to current matrix based methods for protein alignment scoring. It has some unique features, which are well suitable for a range of present and future applications:

(i) *evolutionary independent analysis of sequence similarity* - The scheme is derived without *a priori* knowledge of amino acid distribution or substitution preferences (i.e. independent from biological or evolutionary observations), and thus it may prove particularly beneficial in studies where the evolutionary component is not critical, e.g. protein engineering and design of protein variants;
(ii) *differential conservation of physicochemical properties in alignments* - With its “2-shell” information layers (“inner core” and “external” regions) it was designed to make the customization (i.e. to favor one or more features over the others) of the similarity search powerful yet straightforward; due to its theoretical form, the customization is achievable by direct tuning of the weighting matrix (without any modification of the amino acid description); consequently, aSMM allows the sampling of a broad and diverse search space, to score alignments differentially, depending on (a chosen sets of) different properties; a practical application would be to score alignments with different parameterization schemes: for example, first scoring with the default scheme; and then with *(i)* a scheme favoring size, i.e. higher side chain atoms weight, and with *(ii)* a scheme favoring functional groups and electrostatics; afterwards, the comparison of the outputs could highlight differentially conserved properties, e.g. preferential preservation of electrostatic properties; with similar experiments, the user could be able to detect particular trends (conservation of some properties, over others) in protein alignments;
(iii) *customization and extension to non standard amino acids* - The description of the amino acids can be enriched with new features; indeed, in respect to the basic scheme (Fig 1), some small variants were introduced for Cys, Pro (Fig. 2) in our default scheme (see discussion in the Method section). Similarly, modifications can be introduced to account for non standard amino acids; a simple example could be the most common, naturally occurring, non standard amino acid, selenocysteine; this residue can easily be handled within the aSMM framework (e.g. by representing it as a sort of “super cysteine”, with selenium scoring in the first row, 4th position, with a “higher than sulfur” weight; and the low pKa - around 5, compared to a reference value of around 8 for Cys - reflected in the first column, last position, with a higher fractional weight, e.g. 0.75); in general, many modified amino acids with an overall conserved structure (i.e. similar to a natural type), such as Selenocysteine or Acetyl Phenylalanine, can be treated within the current aSMM framework with few, appropriate modifications;
(iv) *a fundamental step toward Computer Vision based applications* - aSMM was designed specifically to be visually informative, for future utilization with computer vision; indeed, it was developed with this future application specifically in mind (for example, by using specific color codes, data not shown), whereas the sequence and/or the alignments could be read and interpreted automatically (e.g. by an artificially intelligent, computer-vision based, sequence analyzer); the advantages over the simplest analytical approach (i.e. employed here) could be, in time, a winning feature of the Amino Acid Stylization approach presented here.

Image recognition and amino acid stylization can also pave the way to an evolved scoring system, that is sensitive to the surroundings (rather than strictly position specific) and where similarities are scored beyond the isolated value for the *i-th* position, so that to provide a more “fluid” evaluation of protein similarities and of protein/sequence properties.

The work presented is an introduction and a first description of a new approach for protein sequence similarity scoring; it describes a first step into AA stylization approaches, that could evolve in alternative schemes (i.e. also with significant differences from schemes in Fig. 1 and Figure 2), while maintaining the general approach described here. A particular aim of this author is to develop it further as an innovative tool for synthetic biology and protein design, with *(i)* improved representations (that can accommodate some relevant “non standard” features) for non canonical amino acids and *(ii)* with employment of computer vision (i.e. with *ad hoc* color codes for different features) and artificial intelligence to enhance and extend the reach of the similarity search.

## Appendix

Below are the sequences of the pfam alignments used in *Example 1* (note: the employed sequences are a subset of the full pfam alignments, i.e. since the full alignments can be very large and with long stretches of gaps, which would be detrimental to this type of analysis); different families are separated by a # symbol, followed by pfam ID..

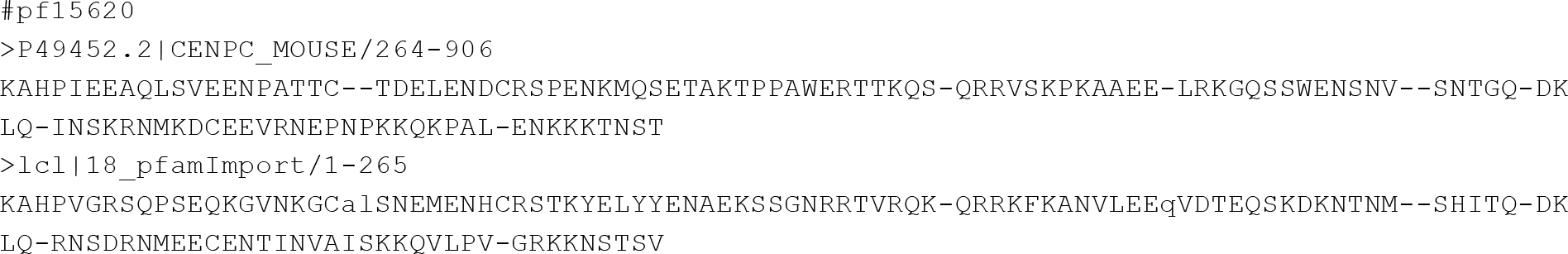

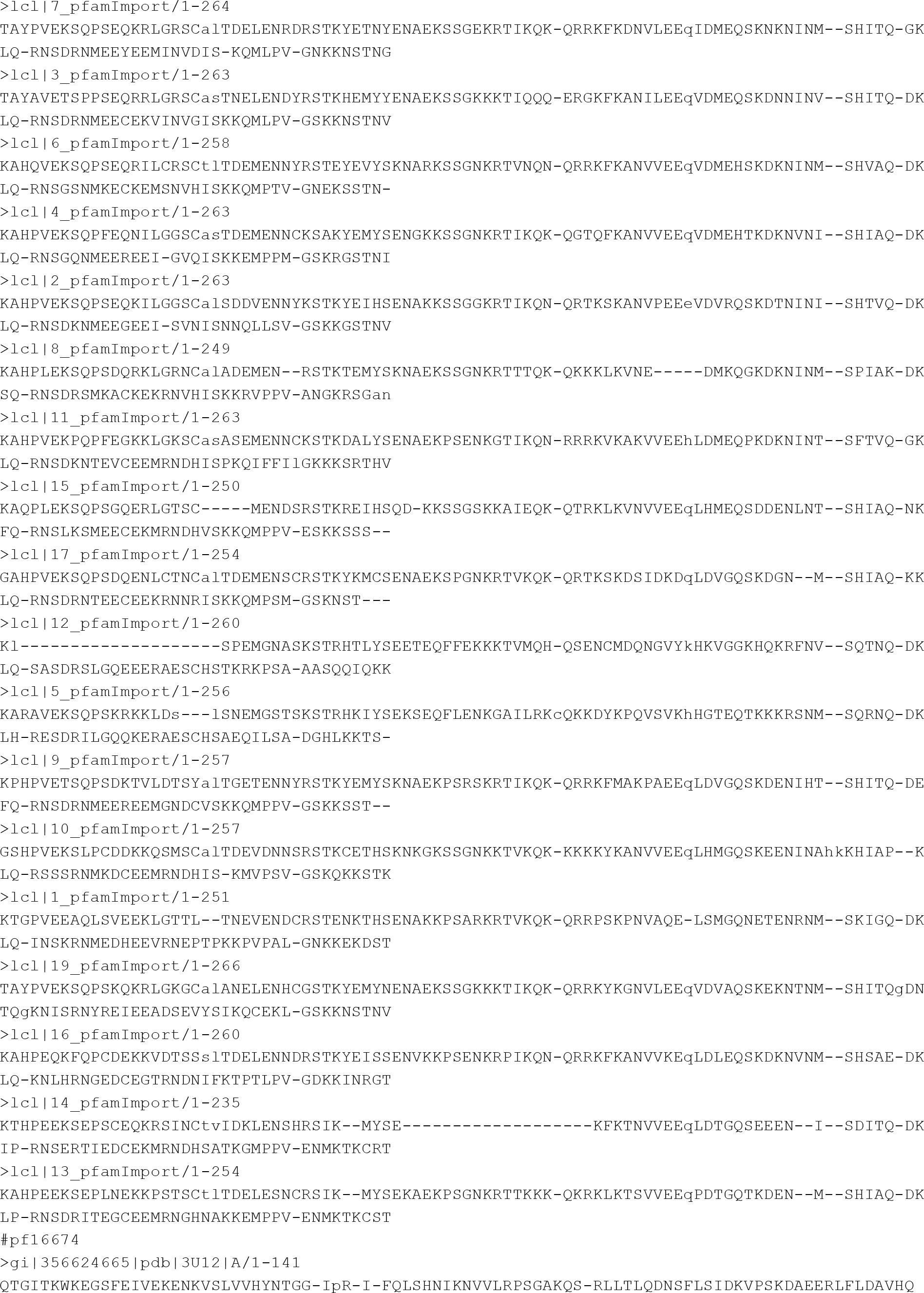

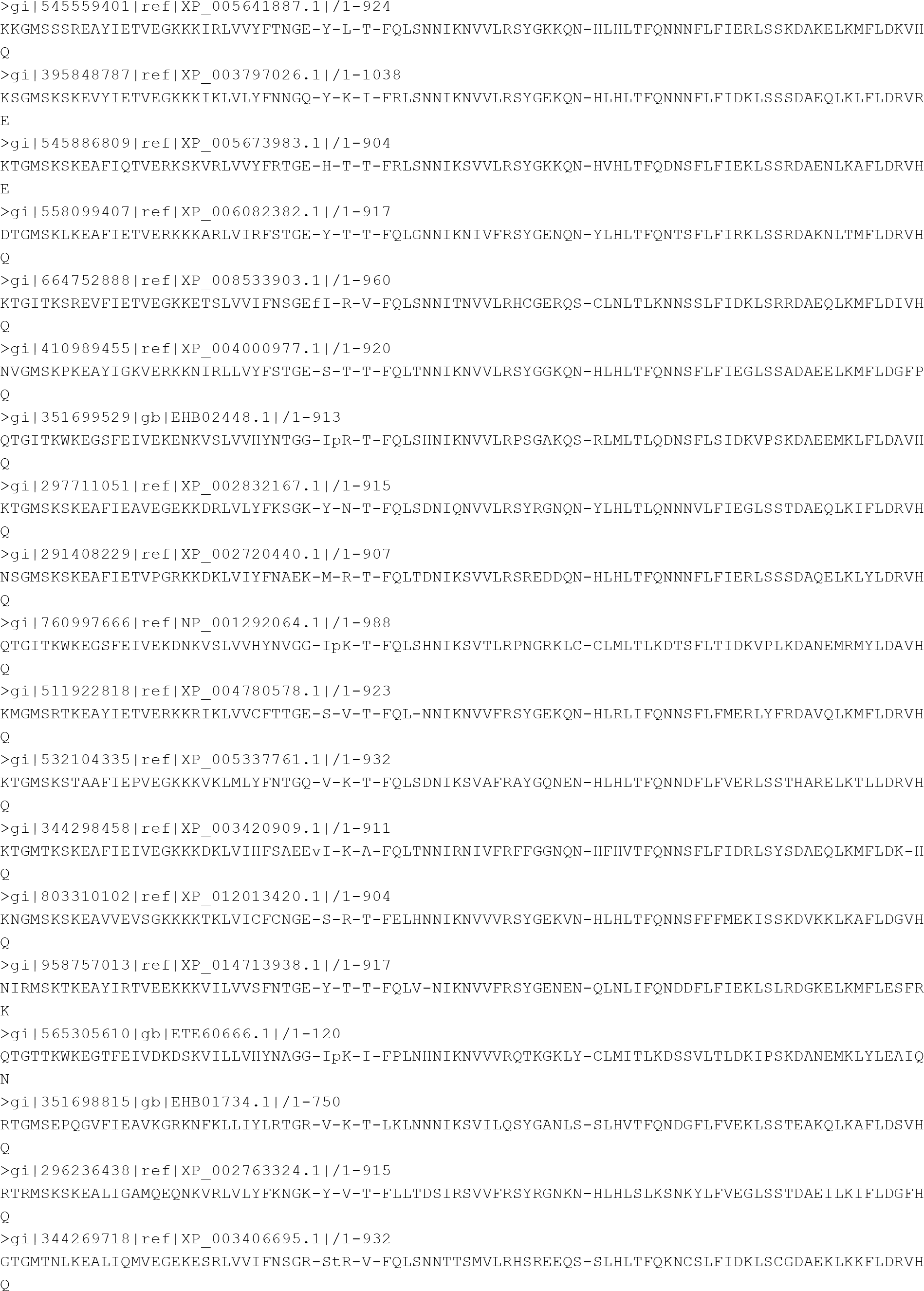

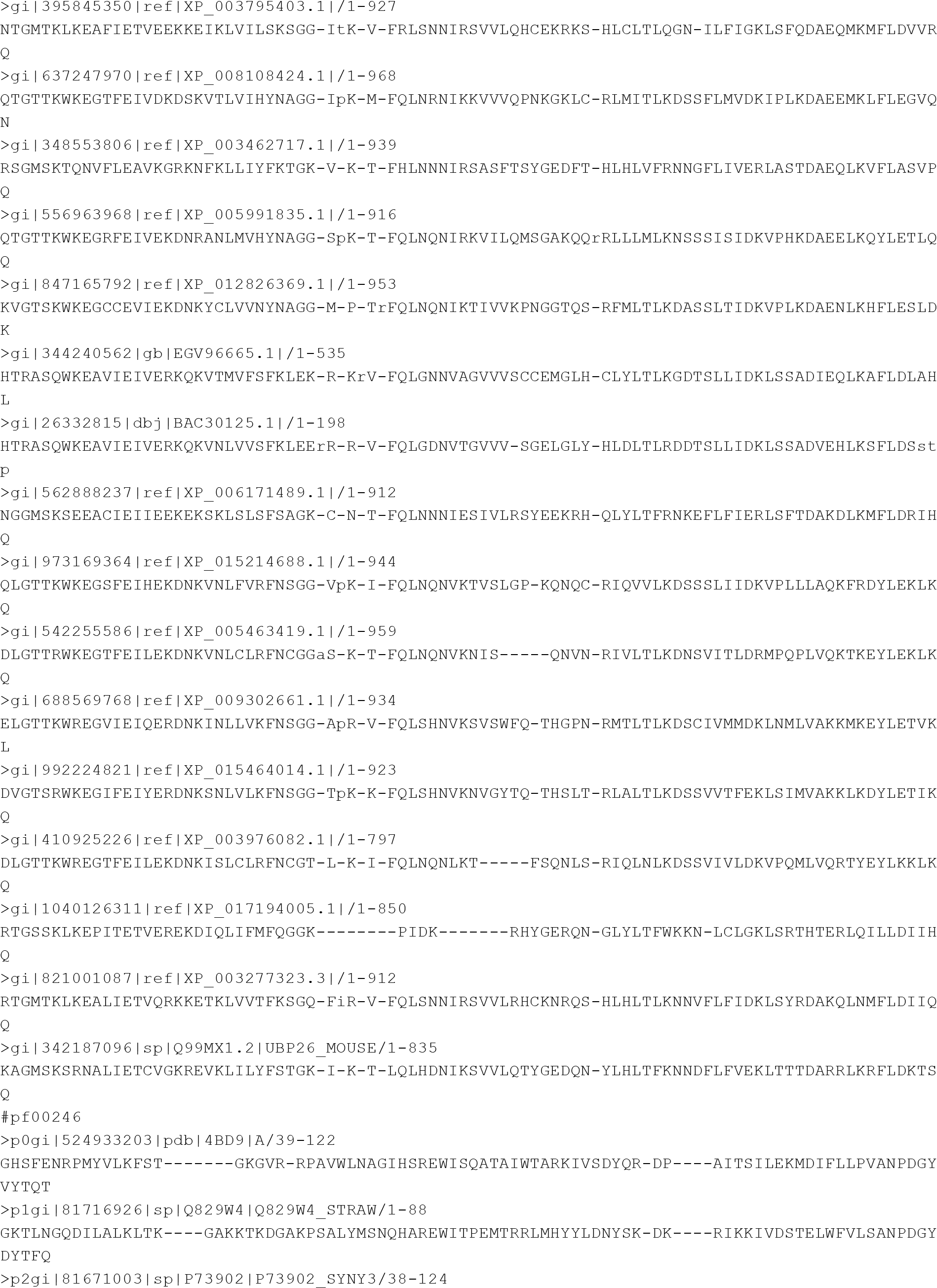

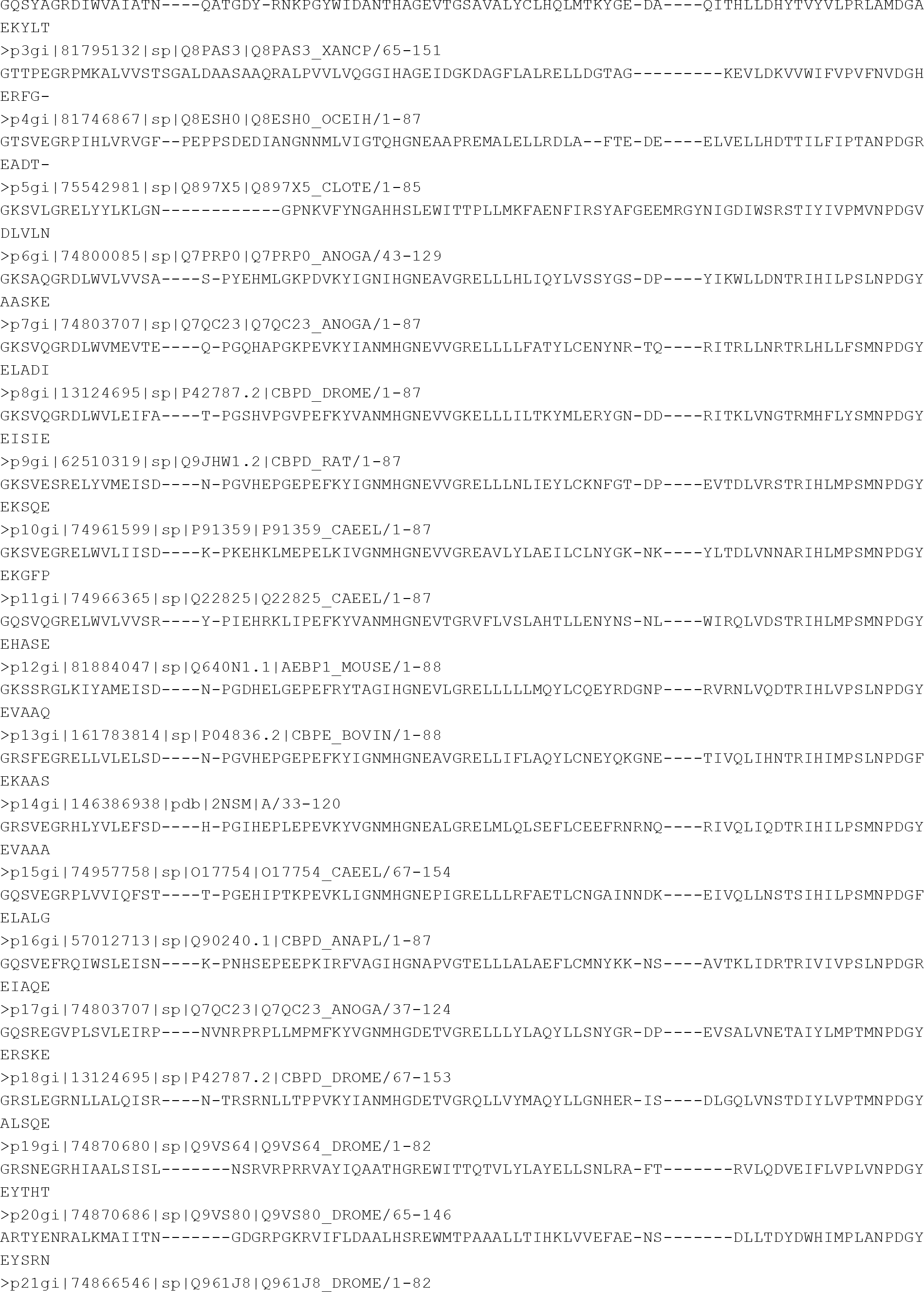

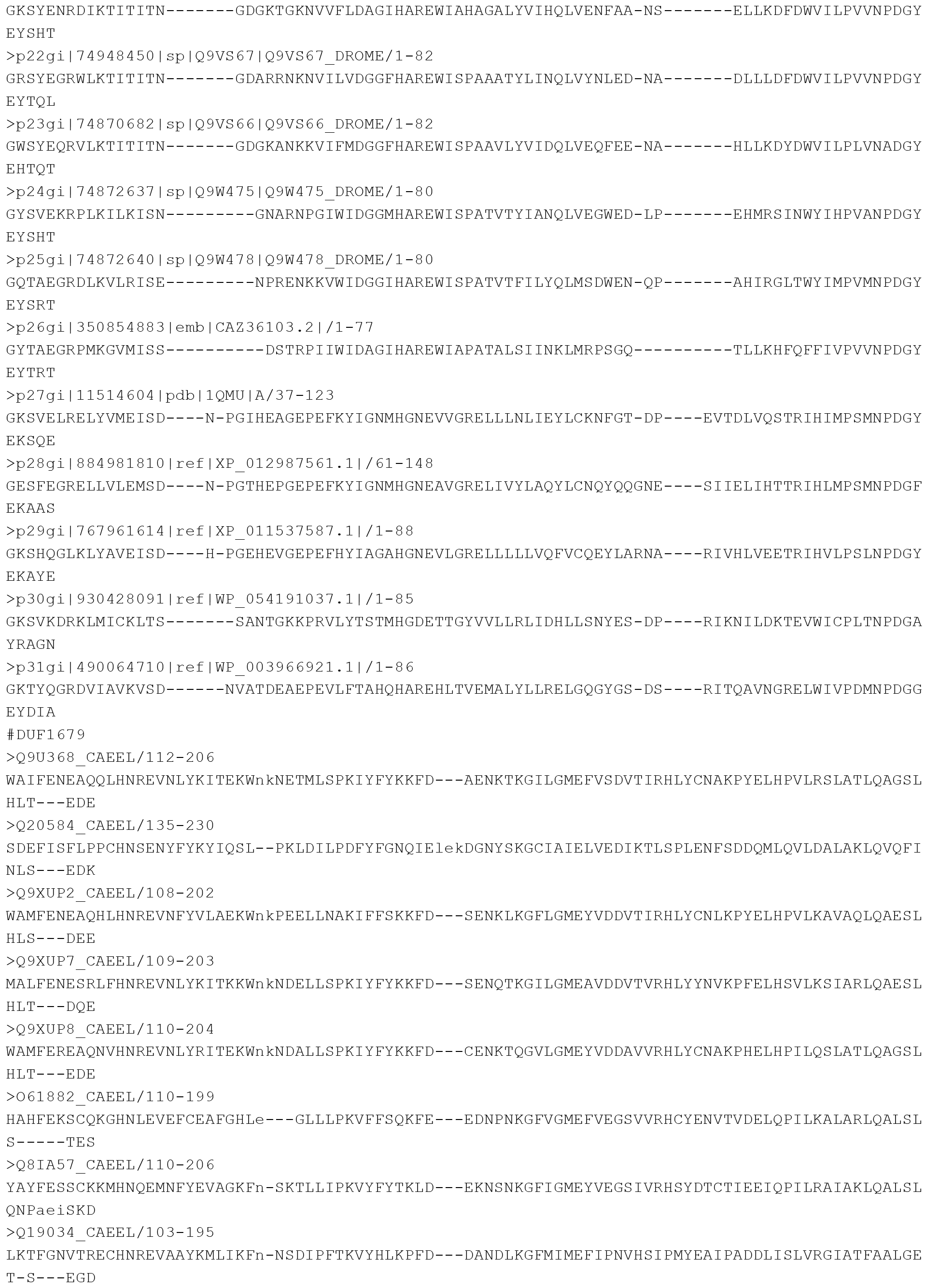

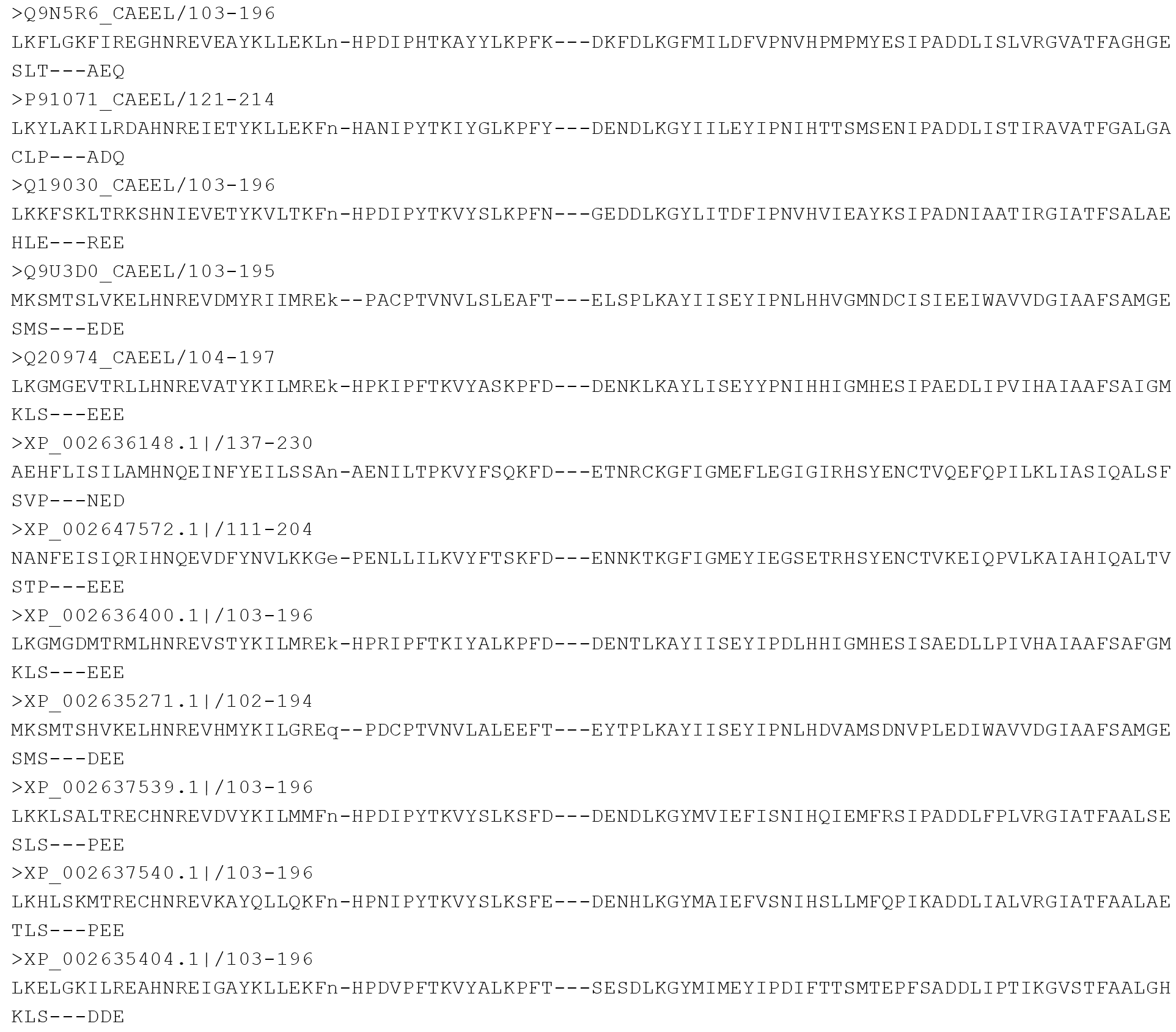

## References

1. Pearson WR. Selecting the Right Similarity-Scoring Matrix. Curr Protoc Bioinformatics. 2013;43:3.5.1–9.

2. Blast: https://blast.ncbi.nlm.nih.gov/Blast.cgi

3. Altschul SF, Gish, W, Miller, W, Myers, EW, Lipman DJ. Basic local alignment search tool. J Mol Biol. 1990 Oct 5;215:(3).403–10.

4. Lipman DJ, Pearson WR. Rapid and sensitive protein similarity searches. Science. 1985 Mar 22;227:(4693):1435–41.

5. Henikoff S, Henikoff JG. Amino acid substitution matrices from protein blocks. Proc Natl Acad Sci U S A. 1992 Nov 15;89:(22):10915–9

6. Styczynski MP, Jensen KL, Rigoutsos I, Stephanopoulos G. BLOSUM62 miscalculations improve search performance. Nat Biotechnol. 2008 Mar;26:(3):274–5.

7. Marino SM, Gladyshev VN. Cysteine function governs its conservation and degeneration and restricts its utilization on protein surfaces. J Mol Biol. 2010 Dec 17;404:(5):902–16

8. Pfam:http://pfam.xfam.org/

